# The Geometry of ATG-Walks of the Omicron SARS CoV-2 Virus RNAs

**DOI:** 10.1101/2021.12.20.473613

**Authors:** Guennadi A. Kouzaev

**Affiliations:** Department of Electronic Systems, Norwegian University of Science and Technology - NTNU, Trondheim, Norway, NO-7491

**Keywords:** Omicron SARS CoV-2 virus, genomic ATG-walk, RNA fractal analysis

## Abstract

In this message, the complete RNA sequences (GISAID) of Omicron (BA.1 and BA.2) SARS CoV-2 viruses are studied using the genomic ATG-walks. These walks are compared visually and numerically with a reference RNA (Wuhan, China, 2020), and the deviation levels are estimated. Statistical characteristics of these distributions are compared, including the fractal dimension values of coding-word length distributions. Most of the 17 RNA ATG walks studied here show relatively small deviations of their characteristics and resistance to forming a new virus family.

## 1. Introduction

At this moment, the attention of biomedical specialists is turned to a new mutation of the SARS CoV-2 virus discovered in Botswana, South Africa, and Hong Kong and denoted by the World Health Organization (WHO) as Omicron [1] or BA1,2 according to the PANGO classification system [2],[3]. It is distinguished from other virus lines by extensive mutation and fast distribution worldwide. In a few weeks, the virus is registered in many tens of countries, and, according to the first reports, it shows increased contagiousness. The origin of this lineage is not known precisely. It is supposed that it arose in a person with a poor immune system co-infected by several virus-triggered diseases [4],[5]. The resistance of vaccines towards this virus is under the studies today.

One of the methods of investigation of RNAs is the numerical mapping of genomic sequences and their following processing by different techniques [6]–[12]. The results of these mappings can be shown graphically. For instance, the positions of each of four nucleotides in a sequence form a trace in 4-D space called a genomic walk [13]. There are many ways to show these walks in the 2-D or 3-D spaces. Each of them has its property preferable for specific analyses. The repeating-pattern walks can be used if a DNA sequence is too long for visual studies [8].

One of the repeating patterns in genomes is an ATG triplet that starts each codon, and the ATG-walks are proposed in Refs. [14],[15] for qualitative and quantitative analyses of virus RNAs. It is shown that for the relatively stable coronaviruses like SARS CoV-2 and MERS ones, the ATG-curves are tightly placed close to each other. The unstable and dangerous viruses like Dengue and Ebola are described by a relatively strong divergence of their ATG-curves expressed in numerical values.

In this short message, the available complete genomes of the Omicron virus [16] are illustrated by ATG-curves. Their properties are compared with each other and those of a Wuhan’s registered SARS CoV-2 virus (GenBank [17]).

## 2. Methods

The pattern search method is described in [14], where the binary symbols represent the characters of RNA and pattern query (base) of an arbitrary length to accelerate the search. Then, the Hamming distance value between binary-expressed symbols in the sequence and base is calculated. The identical symbols give zero value of the calculated distance. This operation is fulfilled each step along the studied RNA sequence.

A string of zeros corresponding to the query is considered a place of the searched pattern. Then, the position of the first base symbol in the sequence is calculated, memorized, and plotted, and it is the pattern coordinate. A trajectory is built to depend on the consequence number *y_i_* of patterns on their coordinates *x_i_* in the studied series of nucleotides. For visual convenience, these points are connected by lines. These lines make a trajectory of an imaginable walker. If ATG-triplet is chosen as a pattern, this trajectory is called an ATG-walk [14].

Additionally to ATG pattern walk illustration and analysis, the fractal dimension of the word-length distributions are studied following the Refs. [14],[15]. For this purpose, the FracLab software tool [18] is used. As a rule, the viruses of the same family have fractal dimension values close to each other.

## 3. Results

In this message, the ATG-walks of the Omicron SARS CoV-2 are investigated and analysed in comparison with a reference virus MN988668.1 (GenBank [17]) registered in Wuhan, China. The numerical results of the study are consolidated in Table 1, Appendix 1, where the number of ATG-triplets in the modeled sequences, word median length, RMS word length, and fractal dimension of word-length distribution are given for 17 RNAs arbitrary chosen from the GISAID database [16]. The RNA word definition is introduced in [14], and it is an ATG-triplet plus codon. The word length is given in nucleotides. Only the sequences without missed nucleotides (marked by N in GISAID) are used for the study. In Fig. 1, the 18 mentioned ATG-curves are shown for comparison, which numbers correspond to the ones in Table 1, Appendix 1.

**Fig. 1.**
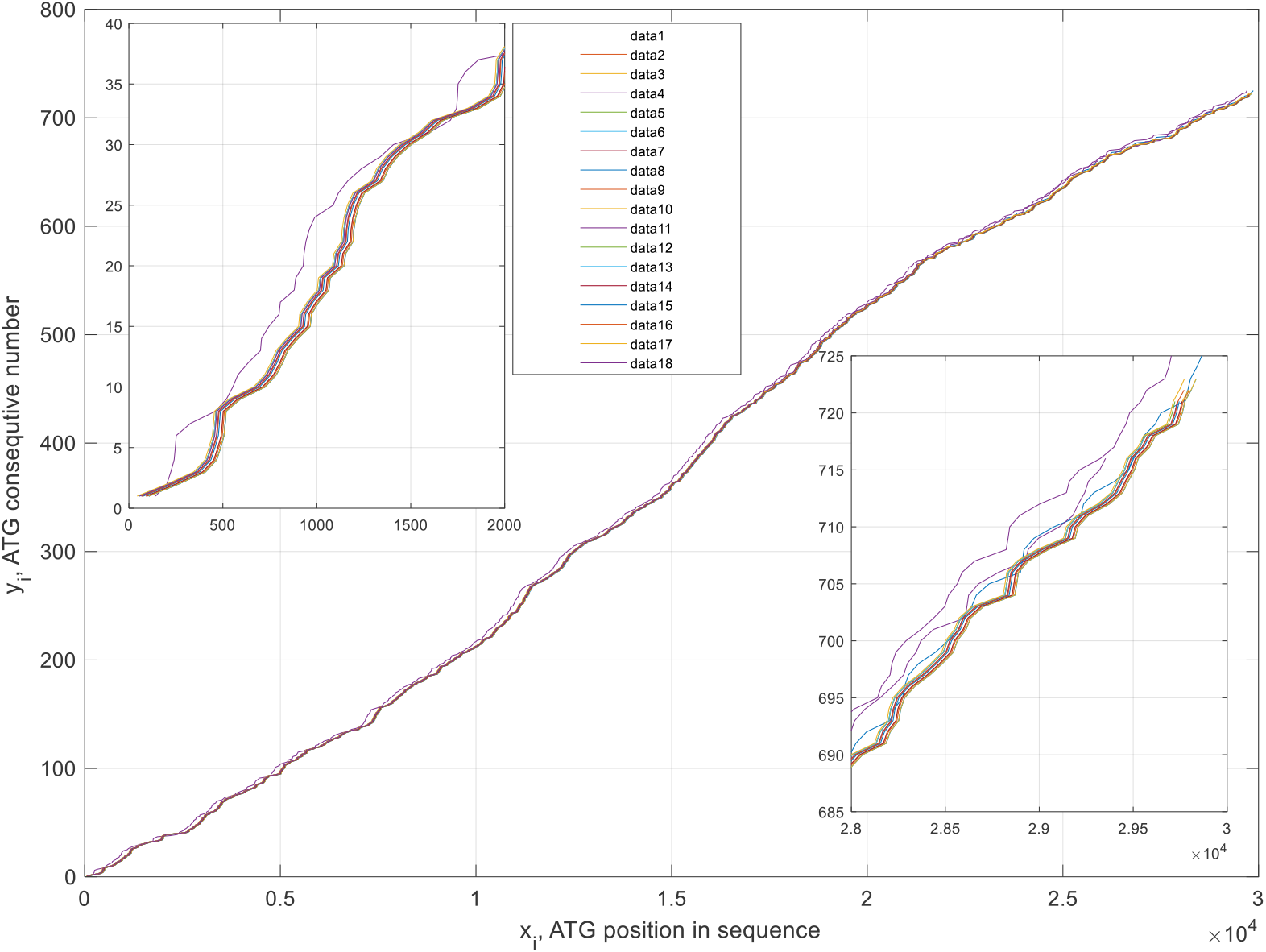
Distributions of ATG-triplets of nineteen SARS Cov-2 complete RNA sequences (rows 1-18, Table 1, Appendix 1). Inlets show the ATG-distributions at the beginning and the end of genome sequences.

It is seen that the walks are geometrically close to each other and to the genomic walk of the reference virus, i.e., the Omicron virus has not been transformed into a new family. Deviation of the curves is estimated in the order 1-2%. Its genetic peculiarities and virus coat protein shapes have been classed recently to a new BA.1 and BA.2 lineages [2],[3]. Some attention should be paid to the RNA from Australia (row 4, Table 1, Appendix 1). It shows a more significant difference than other trajectories (Fig. 1, upper violet curve).

Additionally to the curve plotting and their analyses, the fractal dimension values for the word-length distributions are calculated using the FracLab software tool [18]. It is seen from Fig. 2 that these values computed for the studied RNAs are close to each other, excepting an Omicron virus registered in Australia (sample4, Table 1, Appendix 1).

**Fig. 2.**
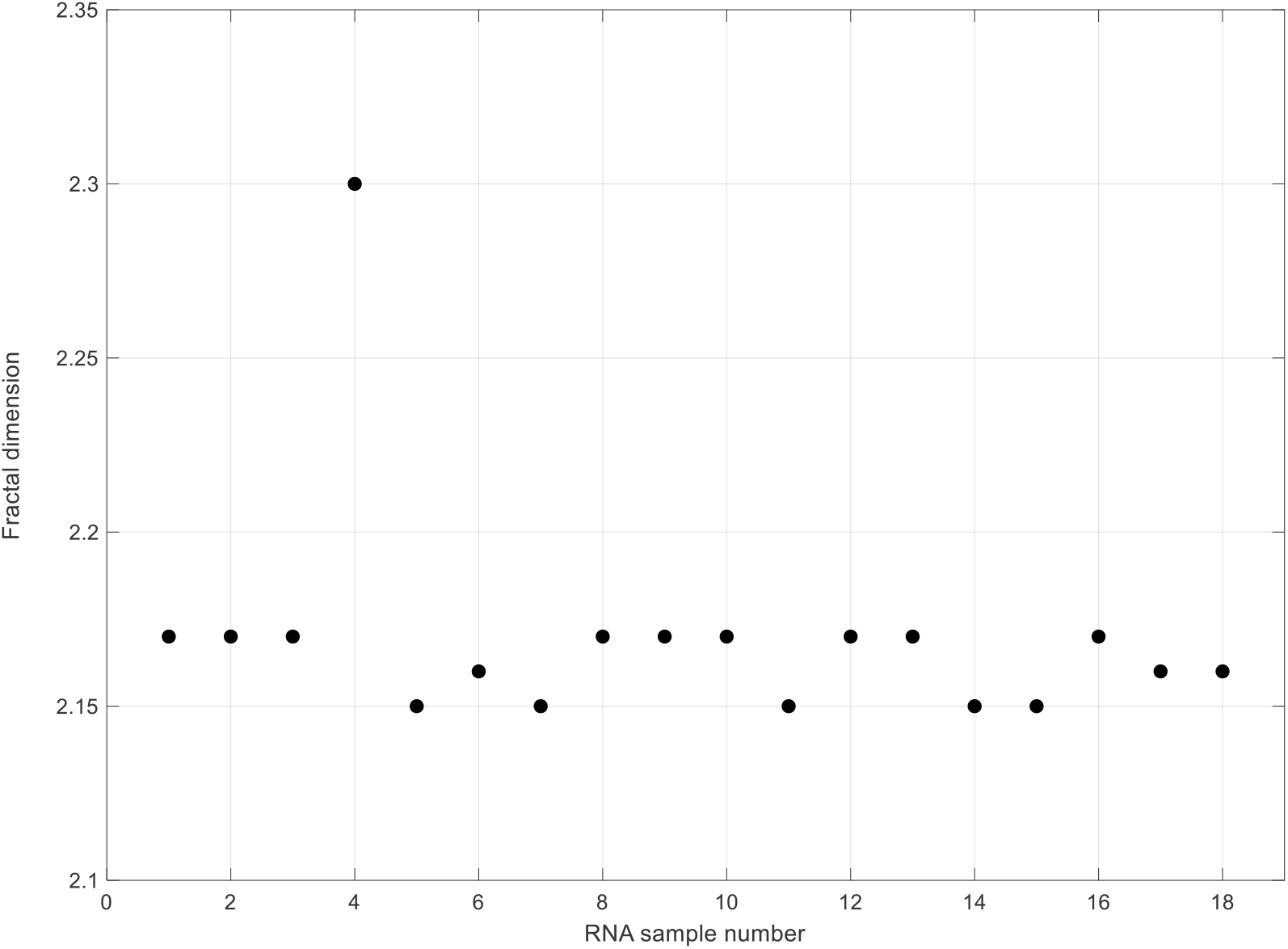
Fractal dimension of the word-length distributions of the complete genome sequences from Table 1, Appendix 1.

Another simple quantitative way to compare the ATG-walks is used in Ref. [13]. We developed a simple algorithm on a detailed comparing of ATG distributions of different virus strains. Each numbered ATG triplet (*y_i_*) has its coordinate *x_i_* along a sequence. Thanks to some mutations, the length of some coding words is varied together with the coordinate *x_i_* of this triplet. Comparing these varied coordinates with a reference sequence allows estimating these mutations. Researching genomic data, we suppose that the error of nucleotide sequencing is negligibly small; otherwise, the results of comparisons would be highly noised.

There are different techniques for comparing sequences known from data analytics, including, for instance, calculation of correlation coefficients of unstructured data sequences, comparisons calculating the data distances, clustering of data, and visual analysis.

In our case, we calculate the absolute difference between the coordinates of ATGs of the same consecutive numbers in the compared sequences. This operation is fulfilled only for the sequences of the same number of ATG triplets; otherwise, excessive coding words are neglected in comparisons. Of course, such a technique for comparing geometrical data has its disadvantages. It means, if a compared sequence has several ATG triplets less than the number of ATG one in a reference sequence, the ATGs of reference RNA are excluded from comparisons. Still, it allows, anyway, to obtain some information on mutations of viruses in a straightforward and resultative way that will be seen below.

This approach supposes choosing a reference nucleotide sequence to compare the genomic virus data of other samples. Here, it is a complete genomic sequence MN988668.1 (GenBank [17]) submitted from Wuhan, China (row 1, Table 1, Appendix 1).

The ordinate axis Δ*x* in these plots shows the deviation of the coordinates *x_i_* of ATG triplets from the ATG coordinates of the reference sequence of the same consecutive ATG number *y_i_*. As a rule, due to the different number of non-coding nucleotides at the beginning of the sequences, the curves have constant biasing along the Δ*x* axis (Fig. 3).

**Fig. 3.**
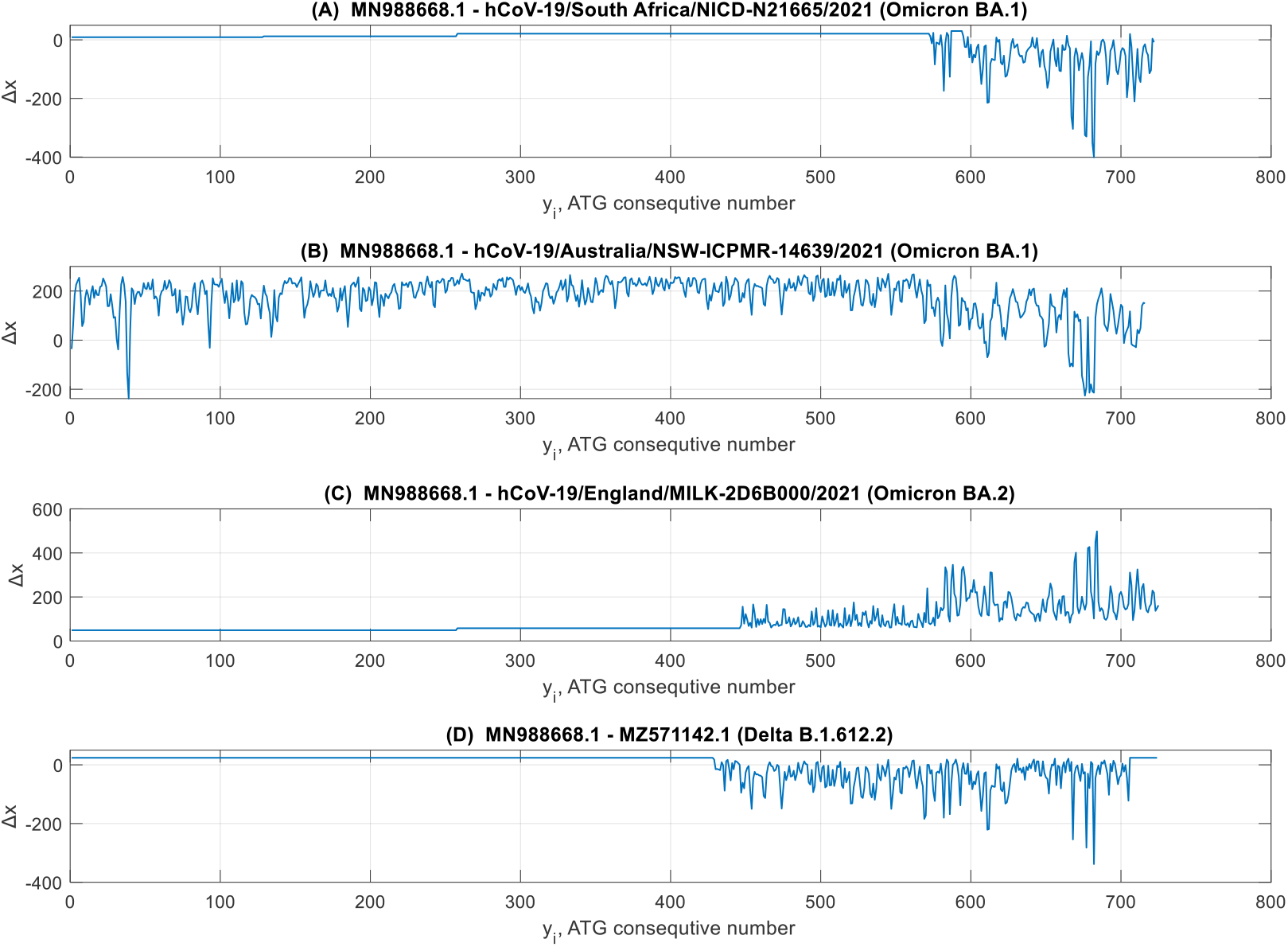
Comparisons of ATG-walk data for the Omicron (BA.1 and BA.2), Delta (B.1.612.2, GenBank)) and reference MN988668.1 (GenBank) SARS CoV-2 virus RNAs.

Straight parts of these curves mean that the ATG positions of a compared sequence are not perturbed regarding the corresponding coordinates in the reference RNA. In these parts, there are no mutations, or the modifications are only with the variation of coding word contents without affecting the codon’s lengths.

A typical deviation curve is shown in Fig. 3A. Comparison of a BA.2 virus (Fig. 3B) provides another shape, but more data is needed to conclude if there is any coupling of these shapes to the lineages of the compared viruses.

Some attention should be paid to a virus sample from Australia (row 4, Table 1, Appendix 1). It has a trajectory with the most significant deviation from the reference trajectory (See Fig. 1, violet curve numbered by 4). Its fractal dimension is twice larger than the typical value for SARS CoV-2 RNA’s samples. At the same time, the calculated difference curve (Fig. 3B) deviates from those for Omicron BA.1 RNAs (Fig. 3A). For comparison, Fig. 3D is a difference curve given for a Delta virus [15].

The next set of curves is given to compare a BA.2 virus with some others, including BA.1 and Delta virus RNAs. Analysing Fig. 1 and Figs. 3 and 4, it is seen that the walks start differentiation at the sequence areas with nucleotides numbered by 23000 (Fig. 3A) and 15000 (others in Figs. 3 and 4). An exclusion is with the virus hCoV-19/Australia/NSW-ICPMR-14639/2021, which is completely different from others. In general, the RNA’s geometrical perturbations Δ*x* do not exceed 2%. In spite of these relatively small distortions, their biological consequences can be strong enough, considering, for instance, the Delta and Omicron pandemic waves in the world.

**Fig. 4.**
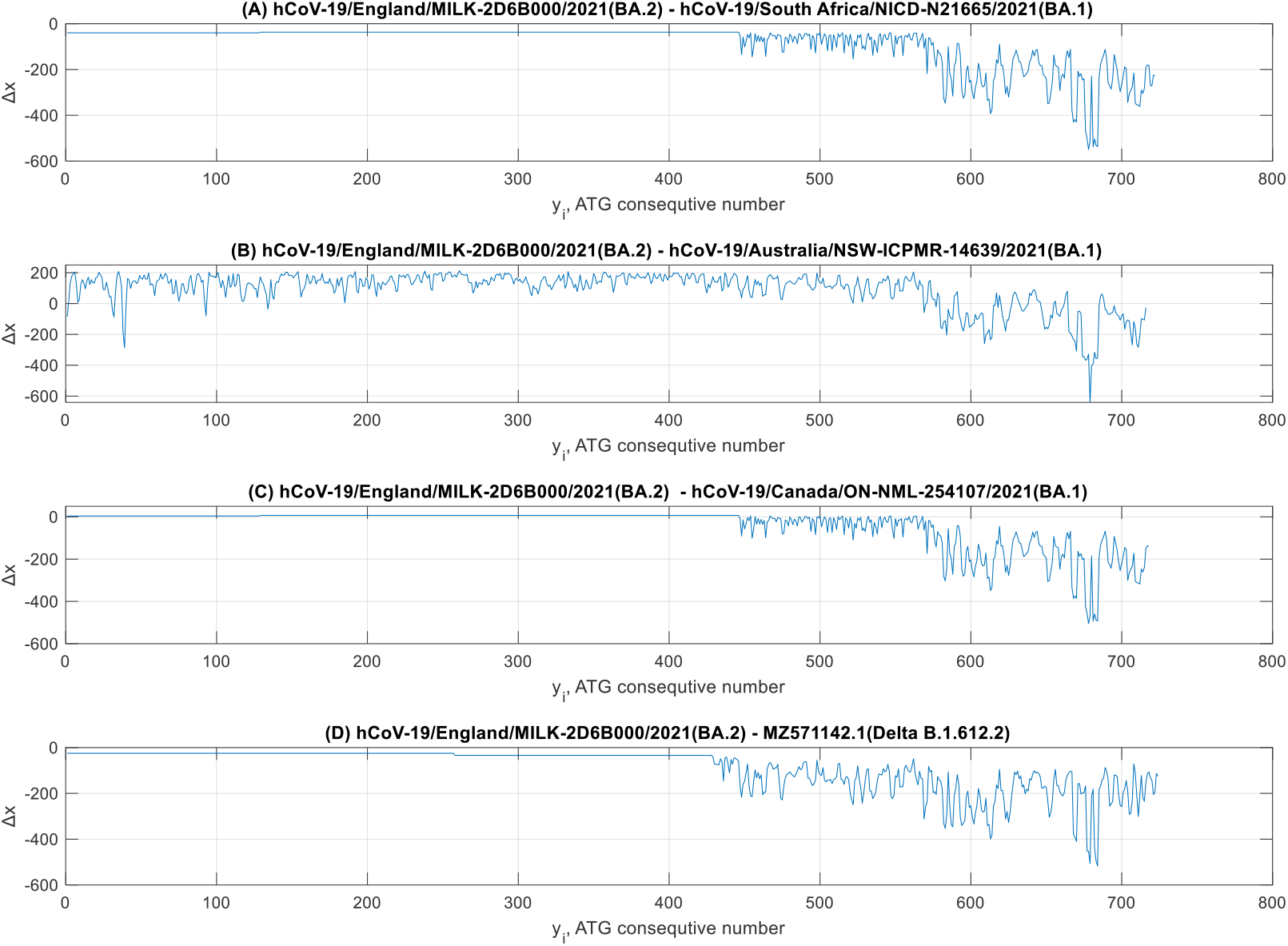
Comparisons of ATG-walk data for the Omicron (BA.1), Delta (B.1.612.2, GenBank)) and reference hCoV-19/England/MILK-2D6B000/2021 SARS CoV-2 virus RNAs.

## 4. Conclusion

In this message, the ATG-pattern walks have been calculated and analysed for the Omicron SARS CoV-2 virus RNAs registered at the GISAID database [16] using the calculation technique from [14],[15].

The ATG triplets are the start-up ones of codons, and their genomic trajectories can be considered the schemes of RNAs having relative stability. At the same time, some mutations change the codon length, and they can be ‘felt’ by ATG-trajectories. Then, ATG-walk method proposed in Ref. [14] is a convenient tool for geometrical mapping of the mutations varying the codon length.

In this way, more than 17 RNAs of the BA.1 and BA.2 Omicron SARS CoV-2 virus samples [16] have been studied. The ATG-trajectories of these RNA samples are close to each other in the limits of 1-2% deviation. Statistical studies of these samples (Table 1, Appendix 1, and Fig. 2) confirm the proximity of almost all of them.

Additionally to visual analyses of the ATG-trajectories, the coordinates of ATG-triplets in sequences have been compared numerically with the reference RNA samples MN988668.1 (GenBank, Wuhan, China, 2020) and hCoV-19/England/MILK-2D6B000/2021 (BA.2 Omicron). The detailed distribution of the deviation of the coordinates regarding the reference samples confirms the above-mentioned difference estimates in the limits of 1-2%. At the same time, one of the samples of RNAs registered in Australia (hCoV-19/Australia/NSW-ICPMR-14639/2021) shows increased deviation of its ATG-trajectories and their fractal dimension values regarding the used reference RNAs (Figs. 1-4). It may require additional attention to control the distribution of viruses of this type of RNAs if such samples would be found further.

## Acknowledgments

The author thanks the GISAID [16] and GenBank^®^[17] and genetic data banks, and all researchers placed their genomic sequences in them. The online text processing service of https://onlinetexttools.com/ is appreciated.

## Funding

Not applicable

## Declarations

### Ethical approval and consent to participate

Not applicable

### Consent for publication

Not applicable

### Competing interests

The author declares that they have no conflicts of interest that are relevant to this research paper.

## Appendix 1

Results of statistical characterization of complete genetic sequences of the Omicron and reference (Wuhan) SARS CoV-2 viruses

**Table 1.**
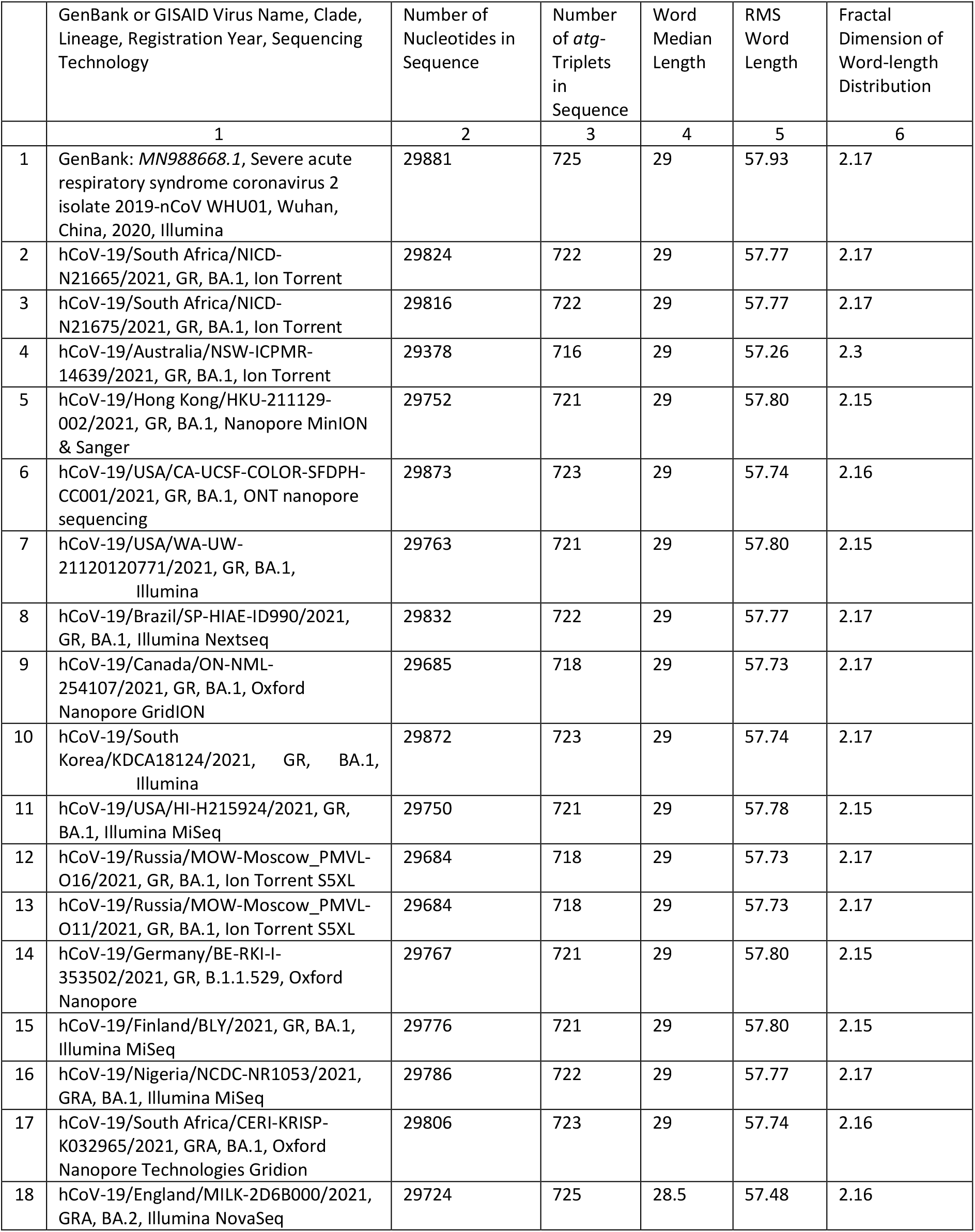
Severe acute respiratory syndrome coronavirus 2 (SARS Cov-2), *atg*-walks

## Notes

### Competing Interest Statement

The authors have declared no competing interest.

## References

1. COVID-19 Weekly Epidemiological Update, Edition 69, published 7 Dec., 2021, World Health Organization (WHO).

2. PANGO Lineages: Latest epidemiological lineages of SARS-CoV-2 [https://cov-lineages.org/index.html].

3. Á. O’Toole, E. Scher, A, Underwood, et al., Assignment of epidemiological lineages in an emerging pandemic using the pangolin tool, Virus Evolution, 7, 1–9, 2021. https://doi.org/10.1093/ve/veab064

4. E. Callaway, Beyond Omicron: what’s next for COVID’s viral evolution. Nature, 07 Dec., 2021.

5. A. Venkatakrishnan, P. Anand, P. Lenehan, R. Suratekar, B. Raghunathan, M. J.M. Niesen, and V. Soundararajan, (2021, December 3). Omicron variant of SARS-CoV-2 harbors a unique insertion mutation of putative viral or human genomic origin. https://doi.org/10.31219/osf.io/f7txy.

6. G. Meister, RNA Biology: An Introduction. Weinheim, Wiley-VCH, 2011.

7. K.R. Kukurba and S.B. Montgomery, RNA sequencing and analysis. Cold Spring Harb Protoc., 11, 951–967, 2015. https://dx.doi.org/10.1101%2Fpdb.top084970

8. B. Brejová, T. Vinar, and M. Li, Pattern Discovery. In: Krawetz S.A., Womble D.D. (eds) Introduction to Bioinformatics, Humana Press, Totowa, NJ, 2003.

9. P.D. Cristea, Conversation of nucleotides sequences into genomic signals, J. Cell. Mol. Med., 6, 279–303, 2002. https://doi.org/10.1111/j.1582-4934.2002.tb00196.x

10. M. Randic, M. Novic, and D. Plavsic. Milestones in graphical bioinformatics. Int. J. Quantum Chem., 113, 2413–2446, 2013. https://doi.org/10.1002/qua.24479

11. P.P. Vaidyanathan, Genomics and proteomics: A signal processing tour. IEEE Circ. Syst. Mag., 4th Quarter, 6–28, 2004. https://doi.org/10.1109/MCAS.2004.1371584

12. J.V. Lorenzo-Ginori, A. Rodríguez-Fuentes, R.G. Ábalo, R. Grau, and R.S. Rodríguez, Digital signal processing in the analysis of genomic sequences. Current Bioinformatics, 4, 28–40, 2009. https://doi.org/10.2174/157489309787158134

13. J.A. Berger, S.K. Mitra, M. Carli, and A. Neri, Visualization and analysis of DNA sequences using DNA walks. J. Franklin Inst., 341, 37–53, 2004. https://doi.org/10.1016/j.jfranklin.2003.12.002

14. A. Belinsky and G.A. Kouzaev, Quantitative analysis of genomic sequences of virus RNAs using a metric-based algorithm, bioArxiv preprint: bioArxiv 2021.06.17.448868; Europe PMC: PPR: PPR358597. https://doi.org/10.1101/2021.06.17.448868

15. A. Belinsky and G.A. Kouzaev, Geometrical study of virus RNA sequences, bioArxiv preprint: bioRxiv 2021.09.06.459135; https://doi.org/10.1101/2021.09.06.459135; Europe PMC: https://europepmc.org/article/PPR/PPR391263

16. Global Initiative on Sharing All Influenza Data (GISAID) [ https://www.gisaid.org/].

17. GenBank^®^ [ https://www.ncbi.nlm.nih.gov/genbank/].

18. FracLab 2.2. A fractal analysis toolbox for signal and image processing. [https://project.inria.fr/fraclab/]

